# STUDY ON BREEDING PERFORMANCE, SURVIVAL AND GROWTH PERFORMANCE OF COMMON CARP (*Cyprinus carpio* L.) IN FISH FARM OF DHANUSHA DISTRICT, NEPAL

**DOI:** 10.1101/2020.12.29.424661

**Authors:** Eliza Aryal, Madhav Prasad Neupane, Shankar Sapkota, Saroj Shrestha

## Abstract

Though Common carp is one of the most widely cultured species in Nepal, there is insufficient supply of fry and fingerlings of common carp. Therefore, the study was conducted with the objective to observe the breeding performance, survival and growth of common carp. Two consecutive experiments were conducted during February to May 2020. First experiment was conducted in four nursery ponds having same area (30m×17m) to observe the breeding performance and growth of hatchlings. 24 kakabans were kept in each pond and brood females and males were released in each pond in a ratio of 1:2 (female: male) i.e. 7 female and 14 male. Each brood fishes were injected with Luteinizing Hormone Releasing Hormone (LHRH A_2_) @ 2µg/kg body weight for female and @ 1µg/kg body weight to male for induced breeding of common carp. The experimental duration was of 36 days. Second experiment was also conducted in 4 nursery ponds having same area (30m×15m) to observe the survival of fry and the experimental duration was of 50 days. There was no significant difference in water quality parameters (p>0.05) and feeding among the research ponds during the culture period. Water quality parameters were measured on weekly basis. Periodic samplings were done to evaluate the daily weight gain, specific growth rate, survival rate and change in water quality parameters. The average weight gain, daily weight gain and specific growth rate of hatchlings were 0.54 g, 0.02 g/fish/day and 12.16%/day respectively. Similarly the average weight gain, daily weight gain and specific growth rate of fry were 20.43g, 0.41g/fish/day and 7.32%/day respectively. And the average survival rate of common carp fry in fish farm of Dhanusha was 47% at stocking density of 56, 000/ha. Through induced breeding process, the average number of fry produced from 7female having average total weight 19 kg and 14 male having average total weight 28 kg was 1, 38,750.

## 1. INTRODUCTION

### 1.1 Background information

Nepal is rich in water resources. There are more than 6000 rivers and rivulets including big and small. It makes a country with potential for fish farming. Water resources occupied about 3% of the total area of Nepal and about 5, 00, 000 of it may be available for fish farming. Nepal is a land-locked country. Fish production of Nepal is completely relied on inland water resources. Out of total inland water resources, river, lakes and reservoirs comprise 48.8%, paddy fields 49%, swamps around irrigated fields 1.4% and village ponds 0.8% (shrestha, 1999). According to a country profile of Nepal (DOFD, 2004) it was estimated that during 2003/2004 nearly 1, 36, 000 families were involved in aquaculture, fisheries and associated activities. Fish production in Nepal is gradually increasing with a growth rate of 8-9% per year reaching 77, 000mt in 2017, contributing 55, 500 from aquaculture practices and 21, 500 from capture fisheries but this productivity lags far behind from neighboring countries (MOAD, 2017). Pond aquaculture contributes 86.6% in aquaculture production, a carps are the major fishes, which occupied over 95% of total fish production in Nepal (Mishra & Kunwar, 2014).The seven fish species of carps including three Indian major carps such as Rohu (*Labeorohita*), Mrigal (*Cirrhinusmrigala*) and Bhakur (*Catlacatla*) and three Chinese major carps such as Silver carp, Grass carp, and Bighead carp along with common carp.

Although fish farming is our ancient practice, commercial fish farming is fairly a new activity in Nepal. It was initiated in the mid-1940s on a small scale in ponds with indigenous Indian major carp seed from India (FAO, 2016). Further development began in the 1950s with the introduction of the exotic species common carp (*Cyprinuscarpio*) and three exotic Chinese carps, namely Silver carp (*Hypophthalmicthysmolitrix*), Bighead carp (*Aristichthysnobilis*) and Grass carp (*Ctenopharyngdonidella*) in the 1970s (FAO, 2016). The major part of pond fish production takes place southern part of the country-the terai plain.

Dhanusha district, a part of province no.2, is located in the outer terai of Nepal at latitude 26°43’43” North, longitude 85°55’30” East. Its elevation is 74 masl (Wikipedia).Mallaha community (73%) dominated the fish selling business in Dhanusha district (Farheen, Gupta, & Gupta, 2019). In Nepal, annual fish production is 91, 832 Mt. with productivity 4.92 Mt./ha. Similarly, in Dhanusha annual fish production is 5, 502 Mt. with 4.89 Mt. /ha.(CFPCC, 2075/76).

Common carp belongs to the class Osteichthyes (the bony fishes), the order Cypriniformes and the family Cyprinidae. There are two varieties of common carp in Nepal: the scale carp or german carp (*Cyprinuscarpio* var. *communis*) and the mirror carp or Israeli carp (*Cyprinuscarpio* var. *specularis*). Flat body, short and small head, protractile mouth and two pairs of maxillary barbells are its specific features. It’s dorsal fin is long with a sharp spine. Common carp is a bottom feeder, omnivorous. It is one of the most widely cultured species in Nepal. It is the main fish species that contributes to total fish production in the country after Mrigal (29.2%) and it contributes 19.2% to total fish production in the country.(Husen, 2019).

Common carp is a multiple breeder and can breed up to 5 times a year. However, the peak breeding season in Nepal is March/April in terai and April/May in the hills. Sexual maturity attains in the first or second year. It breeds easily in ponds without hypophysation. Artificial breeding with hypophysation is also common. They can breed and spawn at 18-22°C. (V.G Jhingram, 1985).

### 1.2 Problem statement

Fish farm of Dhanusha district has encountered high mortalities of Common carp at hatchling and fry stage. So there is insufficient seed of common carp. Data recording related to growth performance and survival rate of hatchling and fry of common carp is poor in fish farm of Dhansha. Infection, unavailability of suitable diet etc. are some factors responsible for poor survival of fish larvae (Little, Tuan, & Barman, 2002). Not only the fish farm of Dhanusha but many fish hatcheries of Nepal have encountered the same problem. In present scenario, insufficient carp seed has become a major obstacle for the development of fish farming. Lack of proper nursing (caring of one week old hatchlings), poor quality broodstock that results poor quality of eggs and larvae thereby increasing mortality of hatchlings, poor water quality of brood ponds are major causes of poor survival of fish larvae. Majority of hatcheries of Dhanusha are facing problems of asphyxiation, predatory aquatic insects, frogs, water snakes and piscivorous birds like king fish during nursing. (Bhujel & Anal, 2015).Data recording related to growth performance and survival rate of hatchling and fry of common carp is poor in fish farm of Dhanusha.

### 1.3 Rationale

Due to multiple benefits of common carp, its demand is increasing day by day. But its supply is beyond the demand. Similar in case of Dhanusha, there is insufficient supply of common carp fry and fingerlings from fish of Dhanusha. The major reason behind this is low survival rate of common carp larvae. It has become very important to assess the causes behind the poor survival of common carp larvae for increasing the production of common carp. Therefore, this study identifies the major problem regarding the poor survival of common carp fry in Fish farm of Dhanusha. Data related to growth performance of hatchling and fry of common carp in fish farm of Dhanusha can be obtained through this study.

### 1.4 Objectives

#### 1.4.1 General objective

- To study the breeding performance, growth and survival of common carp.

#### 1.4.2 Specific objectives

- To assess the production of hatchling, fry and fingerling of common carp in nursery pond.
- To assess the growth performance of hatchlings and fry of common carp.
- To be acquainted with the problems related to mortality of fry in fish farm of Dhanusha and to find out its survival rate.

### 2. METHODOLOGY

### 2.1 Experimental site and sub-sector

Two consecutive experiments were conducted to assess the breeding performance, survival and growth of Common Carp during February to May 2020 at Fisheries Human Resource Development and Technology Validation Centre, Janakpurdhaam, Dhanusha. It is located in the outer terai of Nepal at latitude 26°43’43” North, longitude 85°55’30” East. Its elevation is 74masl.

**Figure 1.**
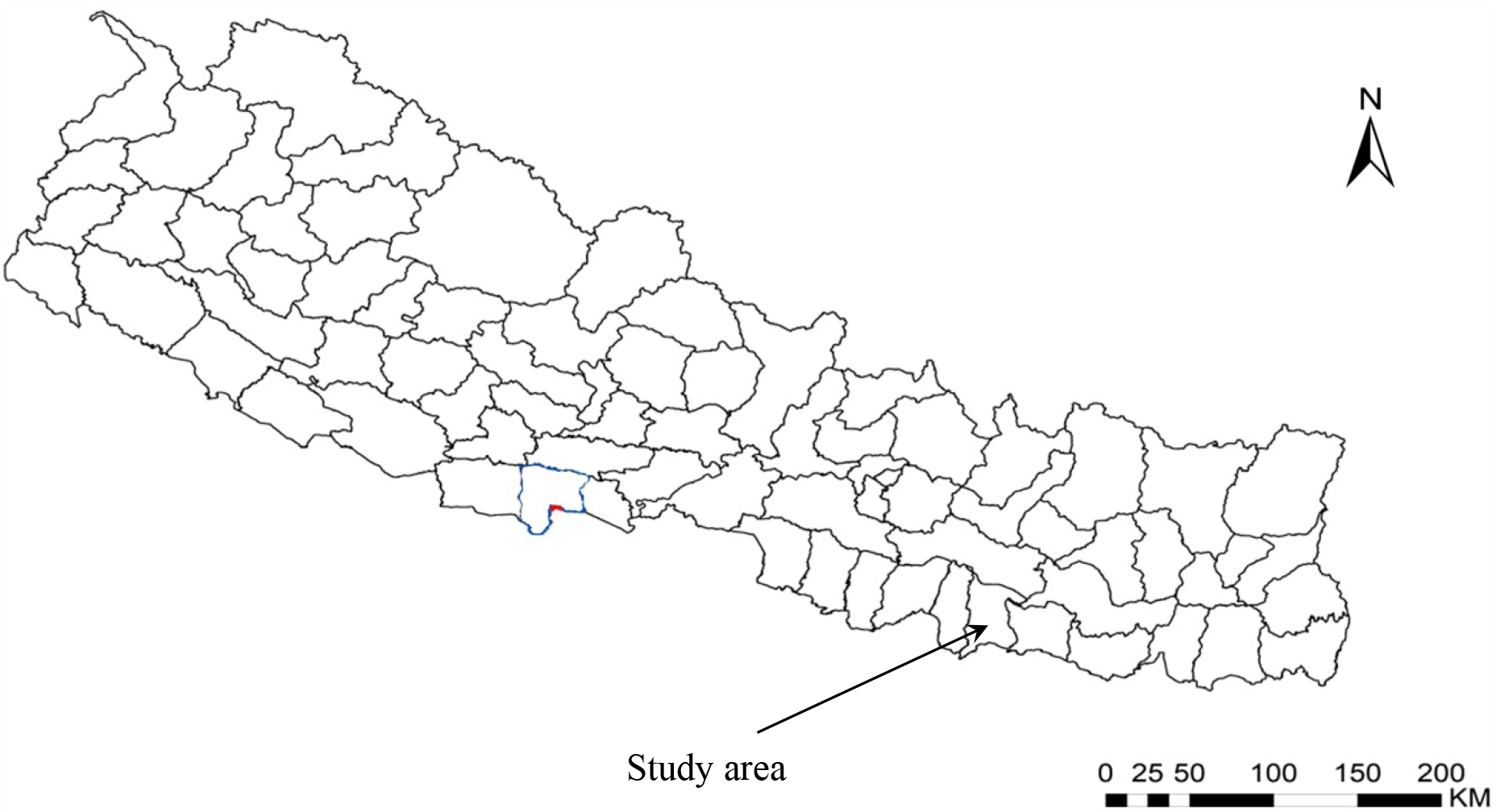
Map of Nepal showing study area.

### 2.2 Details of experiment 1

Experiment 1^st^ was conducted to observe the breeding performance of Common carp and growth performance of its hatchling. Four nursery ponds having same area(30m×17m) were selected for study purpose and were drained. It was conducted from 24^th^ Feb to 30^th^ March so that the experimental duration was 36 days.

#### 2.2.1 Preparation of nursery pond for breeding purpose

Liming was done in all four research ponds @ 20 kg/pond to kill harmful insects, pests and to maintain water P^H^. It was done on 23^rd^ February. On 24^th^ Feb (in the morning) all four nursery ponds were filled with boring water. On 24^th^ Feb (at 4pm) setting was kept in all four research ponds. 24 kakabans were kept in each pond. Kakabans were set at 1.5 ft. above the ground level and along the four sides of rectangular pond. After setting of kakaban water was filled up to 1ft. above the kakaban so that the total depth of water in each pond was 2.5 ft.

#### 2.2.2 Brood source

56 male broods and 28 female broods of common carp were collected from male and female brood ponds of Fish farm, Dhanusha respectively.

#### 2.2.3 Breeding performance

For breeding purpose male and female common carps were harvested and kept in separate brood ponds. On 24^th^ Feb after setting of kakaban, male and female brood fishes were collected. Each brood fish was weighed using electronic weighing machine. Both male and female brood fishes were injected with Luteinizing Hormone Releasing Hormone (LHRH) A_2._Dose for female was 2 microgram/ kg body weight and dose for male was 1 microgram/kg body weight. Injection was given at dorsal fin bases at 45° angle. In each pond brood fishes of Common carp having relatively equal weight were released in the ratio of 1:2(female: male) i.e.7 female and 14 male.

**Table 1.**
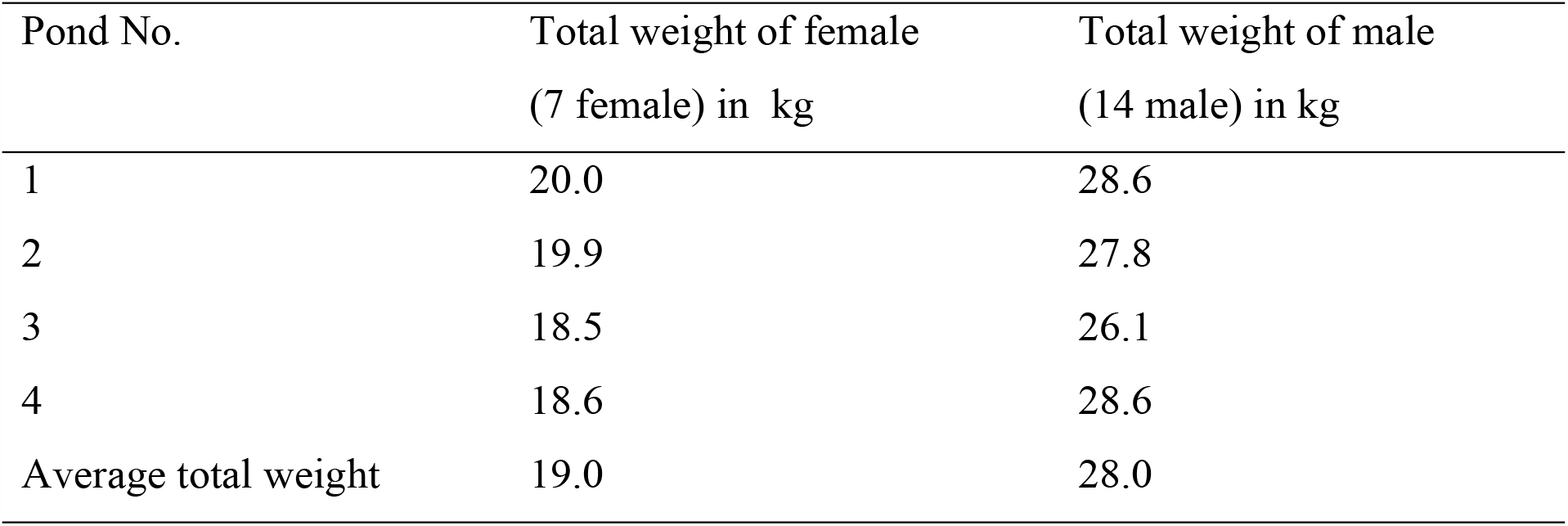
Total weight of male and total weight of female brood fish in each research pond

#### 2.2.4 Harvesting of parent fish after breeding

Brood fish which were kept in pond for breeding purpose were harvested from all ponds on 2^nd^ March after 8 days using net having large hole so that hatchlings can escape from that net. It was done to prevent brood fish from eating their own hatchling and hatchling’s food.

#### 2.2.5 Water quality analysis

Physico-chemical parameters such as water temperature (°C), dissolved oxygen (mg/l) and p^H^ of each pond were measured on weekly basis by thermometer, DO meter and p^H^ meter respectively. Water quality parameters were measured three times per day (6:00 AM, 12:00 AM and 5:00 PM) to know the fluctuation in water quality parameters at different time of a day and their mean values were taken.

#### 2.2.6 Feed and feeding

##### 2.2.6.1 Feeding of hatchling

Feeding of hatchling was started after 3 days of hatching of eggs i.e. 3 days old hatchling. They were fed twice daily between 10-11am and 4-5 pm. Each time 2 eggs were mixed with water and spread thoroughly in the pond. Eggs were given for 1 week. The feeding schedule of all four ponds is same and it is of following type:

**Table 2.**
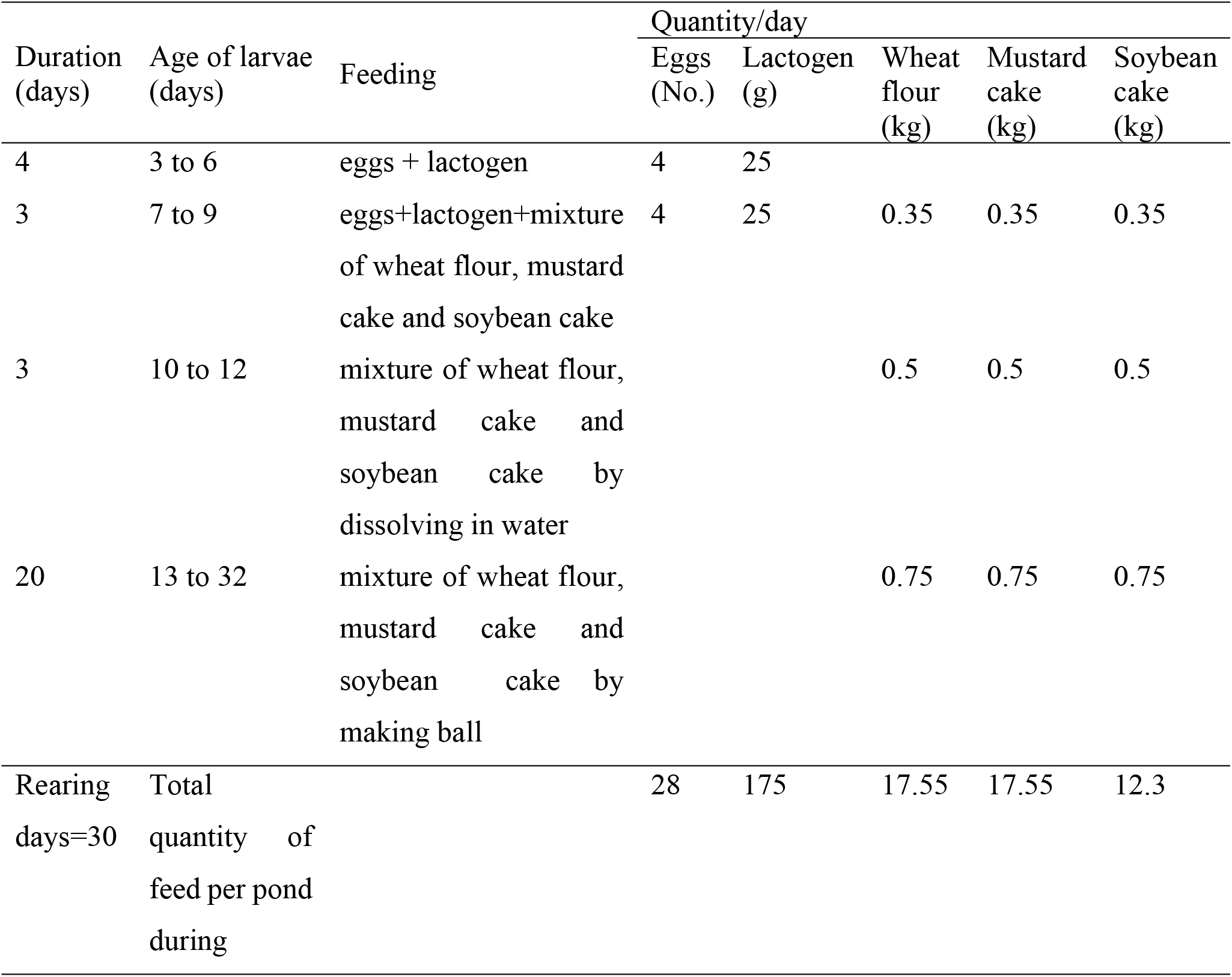
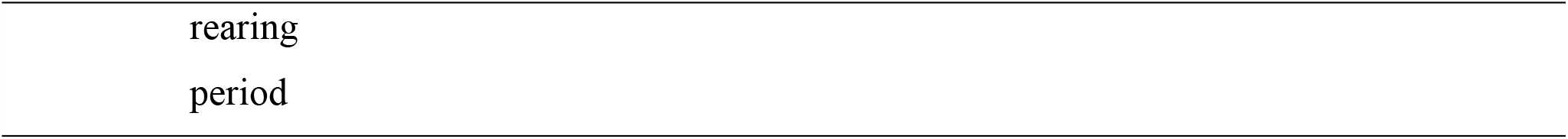
Feeding schedule of hatchling during rearing period of 30 days

| Duration<br>(days) | Age of larvae<br>(days) | Feeding | Quantity/day |  |  |  |  |
| --- | --- | --- | --- | --- | --- | --- | --- |
|  |  |  | Eggs<br>(No.) | Lactogen<br>(g) | Wheat<br>flour<br>(kg) | Mustard<br>cake<br>(kg) | Soybean<br>cake<br>(kg) |
| 4 | 3 to 6 | eggs + lactogen | 4 | 25 |  |  |  |
| 3 | 7 to 9 | eggs+lactogen+mixture<br>of wheat flour, mustard<br>cake and soybean cake | 4 | 25 | 0.35 | 0.35 | 0.35 |
| 3 | 10 to 12 | mixture of wheat flour,<br>mustard cake and<br>soybean cake by<br>dissolving in water |  |  | 0.5 | 0.5 | 0.5 |
| 20 | 13 to 32 | mixture of wheat flour,<br>mustard cake and<br>soybean cake by<br>making ball |  |  | 0.75 | 0.75 | 0.75 |
| Rearing<br>days=30 | Total<br>quantity of<br>feed per pond<br>during |  | 28 | 175 | 17.55 | 17.55 | 12.3 |

##### 2.2.6.2 Feeding of parent fish

Feeding of parent fish was started from27^th^ Feb. 0.5 kg rice bran and 0.5 kg mustard cake were fed per day per pond. Rice bran and mustard cake were mixed with water and fed by making boll.

#### 2.2.7 Harvesting of fry

Final harvesting of fry was done on 30^th^ March after 36 days of setting kakaban. It was done by draining each pond completely. Before completely drying of ponds, fry were netted out as much as possible using a drag net. Total number of fry was counted from each pond and final weight of fry was also measured using microgram weighing balance.

#### 2.2.8 Counting of fry

First of all one plastic bag was filled with water and fry. Fry contained in that plastic bag was counted one by one. And that plastic bag was used as standard for counting fry of all ponds.

### 2.3 Details of experiment

This experiment was conducted on 30^th^ March to observe the survival rate of fry.

#### 2.3.1 Preparation of pond

For this experiment, four nursery ponds having same area (30m×15m) were selected and were drained. Liming was done @ 20 kg/pond.

#### 2.3.2 Stocking of fry

Fry was stocked @56,000/ha in all 4 ponds i.e. 25, 000 fry having mean weight 0.54g were stocked in each pond and cultured for 50 days. Regular monitoring of ponds was done during the culture periods.

#### 2.3.3 Feed and feeding

Wheat flour, soybean cake, mustard cake and pellet feed were mixed in equal proportion. Water was added in that mixture so that feed can be fed to fry easily by making boll. Four bricks were suspended with the help of rope from four side of pond. Bricks were immersed into water. Feed boll was placed over brick and brick was immersed into pond water. This was done to prevent loss of feed and equal distribution of feed to all fish. Quantity of feed fed was increased when their body size increased. They were fed according to their body weight.

#### 2.3.4 Harvesting of fingerling

On 18^th^ May, final harvesting of fingerling was done and also final weight of fingerling was measured. Total number of fingerling was counted from each pond and survival rate was calculated using formula:

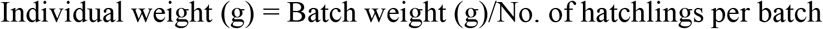

### 2.4. Sampling and sampling technique

To observe the growth performance of hatchling, Simple Random Sampling technique was used. On 2^nd^ March, first sample was taken after 4 days of hatching of eggs (4 days old hatchling) and that weight was considered as initial weight of hatchling. Then weight of hatchlings from each pond was taken on weekly basis up to 30^th^ March (32 days old hatchling) and that weight was considered as the final weight of hatchling. Since weight of single hatchling is negligible and to obtain more precious data, batch weights of hatchling were recorded from each pond using Microgram Weighing Balance. From each pond 5 batches of hatchling were taken as sample per week. Individual weight of hatchling was calculated by using formula:

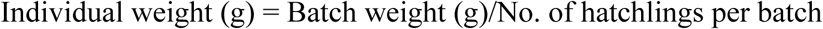

And,

Weight gain, Daily weight gain (g/fish/day) and Specific Growth Rate (SGR) were calculated using the following formula:

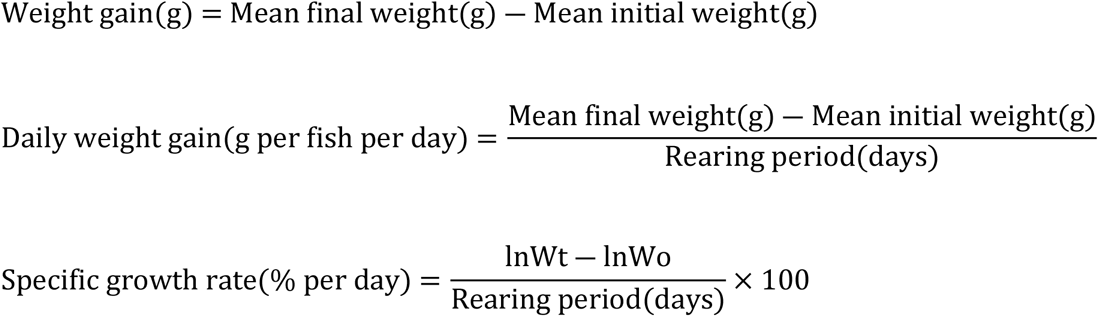

Where, Wt= Mean final weight

Wo= Mean initial weight

### 2.5 Data recorded

Data were collected from research conducted in Fish Farm, Dhanusha. Following types of data were collected during research period:

- Weight of brood fish: Both male and female brood fish used for breeding purpose were weighed using electronic weighing scale.
- Water temperature: Water temperature of all research ponds were measured on weekly basis (three times a day) using thermometer and recorded.
- Water pH: Water pH of all research ponds were measured on weekly basis (three times a day) using pH meter.
- Water DO: Dissolved oxygen of all research ponds were measured on weekly basis (three times a day) using DO meter.
- Weight of hatchling: Weight of hatchlings was recorded on weekly basis using microgram weighing balance.
- Length of hatchling and fry: Length of hatchling, fry and fingerling were taken using scale.
- No. of fry per pond: No. of fry produced from each pond was counted.
- No. of fry stocked: No. of fry stocked in each pond was recorded.
- No. of fingerling produced: No. of fingerling survived from each pond was counted.
- Quantity of feed: Quantity of feed fed during culture periods was recorded.

### 2.6 Data analysis

Data entry and data analysis was carried out using Microsoft Excel software. Single factor ANOVA was also calculated using excel.

## 3. RESULTS AND DISCUSSIONS

All the data collected from the experiment were analyzed to produce fruitful information. Result of the experiment is presented below under different headings.

### 3.1 Breeding performance

Following observations related to breeding performance were made

**Table 3.** Observations related to breeding performance made at different date.

| Date | Observation |
| --- | --- |
| 24<sup>th</sup> Feb-2020 | Setting was kept in all four research pond at about 4 PM. |
| 25<sup>th</sup> Feb-2020 | Eggs laying in kakaban were observed at 4AM in all four research pond after 12 hrs. of keeping set. |
| 28<sup>th</sup> Feb-2020 | Hatchlings were observed in all ponds after 72 hrs. of laying eggs (i.e. after 3 days of laying eggs. |

### 3.2 Production of fry

In pond no.1, 1, 40,000 fry were produced from 7 female and 14 male having total weight 20kg and 28.6 kg respectively. Similarly, in pond no. 2,1,39,000 fry were produced from 7 female and 14 male having total weight 19.9 kg and 27.8 kg respectively. In pond no.3, 1, 37,000 fry were produced from 7 female and 14 male having total weight 18.5 kg female and 26.1 kg male respectively. And 1, 39,000 fry were produced from 7female and 14 male having total weight 18.6 kg and 28.6 kg male respectively in pond no.4. Production of fry was almost same in all four ponds.

On an average, 1, 38, 750 fry of common carp were produced from 7 female having total weight 19 kg and 14 male having total weight 28 kg through induced breeding process.

**Table 4.** Total number of common carp fry produced in different nursery ponds

| Pond No. | Number of fry produced per pond |
| --- | --- |
| 1 | 1,40,000 |
| 2 | 1,39,000 |
| 3 | 1,37,000 |
| 4 | 1, 39, 000 |
| Average | 1, 38, 750 |

### 3.3 Growth performance

#### 3.3.1 Weight gain, daily weight gain and specific growth rate of hatchling

The average initial weight of hatchlings (4 days old hatchling) in Pond1, Pond 2, Pond 3 and Pond 4 were 0.02g, 0.02g, 0.02g and 0.02g respectively. The average final weight gain of hatchlings after 29 days in Pond 1, Pond 2, Pond 3 and Pond 4 were 0.60g, 0.58g, 0.52g and 0.54g respectively. The mean weight gain of hatchlings from pond 1, pond 2, pond 3 and pond 4 were 0.58g, 0.56g, 0.50g and 0.52g respectively. Similarly, the mean daily weight gain of hatchlings from pond 1, pond 2, pond 3 and pond 4 were 0.02g/fish/day,0.02g/fish/day, 0.02g/fish/day and 0.02g/fish/day respectively. And, the specific growth rate (SGR) of hatchlings from pond no. 1, 2, 3&4 were 12.24%/day, 12.20%/day, 12.10%/day and 12.10%/day respectively. The initial weight, final weight, weight gain, daily weight gain and SGR were found almost similar in all four research pond.

Therefore, the average weight gain of hatchlings was 0.56 g. And, the average daily weight gain and SGR of hatchling was 0.02g/fish/day and 12.16 % per day respectively during the rearing period of 29 days.

Body weight of hatchling was increased slowly during the first two week of culture period and rapid increase in body weight was observed after second week. Similar growth pattern was observed in all research ponds. The calculated p-value is 0.994 which is greater than 0.05. That means there was no significant difference in average weight of hatchling among 4 research ponds.

**Table 5.** Initial weight, final weight, weight gain, daily weight gain and specific growth rate of hatchling during the rearing period of 29 days

| Growth parameters | Pond 1 | Pond 2 | Pond 3 | Pond 4 | Average |
| --- | --- | --- | --- | --- | --- |
| Initial weight(g) | 0.02 | 0.02 | 0.02 | 0.02 | 0.02 |
| Final weight(g) | 0.60 | 0.58 | 0.52 | 0.54 | 0.56 |
| Rearing Period(days) | 29 | 29 | 29 | 29 | 29 |
| Weight gain(g) | 0.58 | 0.56 | 0.50 | 0.53 | 0.54 |
| Daily weight gain(g/fish/day) | 0.02 | 0.02 | 0.02 | 0.02 | 0.02 |
| Specific Growth Rate(%/day) | 12.25 | 12.20 | 12.10 | 12.10 | 12.16 |

**Table 6.** Average weight of common carp hatchling at different days after hatching

| Days after hatching | Pond 1(g) | Pond 2(g) | Pond 3 (g) | Pond 4 (g) | Average weight (g) |
| --- | --- | --- | --- | --- | --- |
| 4 | 0.02 | 0.02 | 0.02 | 0.02 | 0.02 |
| 11 | 0.03 | 0.03 | 0.02 | 0.03 | 0.03 |
| 18 | 0.06 | 0.06 | 0.05 | 0.05 | 0.05 |
| 25 | 0.30 | 0.26 | 0.21 | 0.24 | 0.25 |
| 32 | 0.60 | 0.57 | 0.51 | 0.54 | 0.56 |

**Figure 1.**
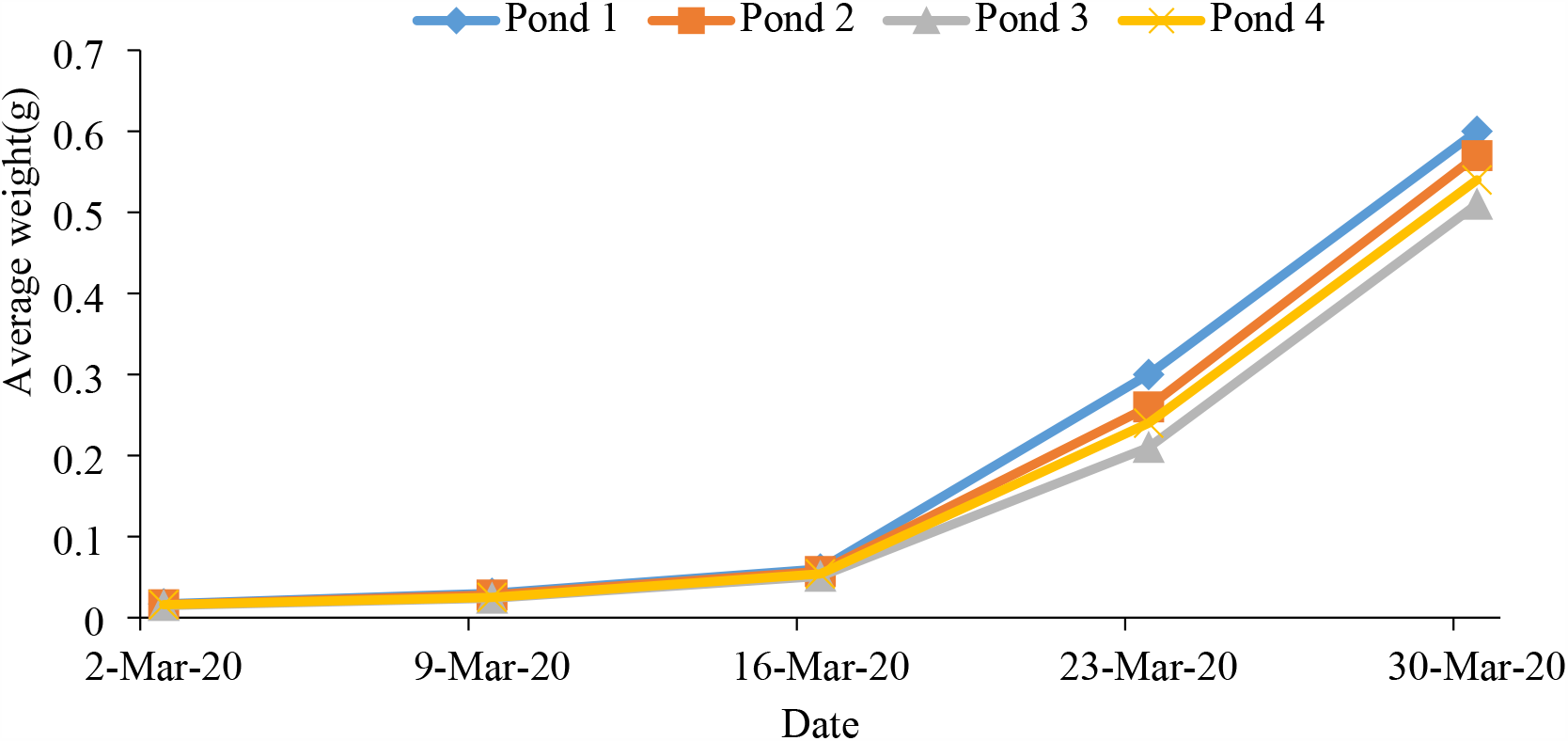
Average weight of hatchling in each sampling.

#### 4.3.2 Final length of hatchling

**Table 7.** Length of common carp hatchling in different ponds after rearing for 32 days

| Sample number | Pond 1<br>(cm) | Pond 2<br>(cm) | Pond 3<br>(cm) | Pond 4<br>(cm) | Average length<br>(cm) | Calculated<br>p-value |
| --- | --- | --- | --- | --- | --- | --- |
| 1 | 2.5 | 2.6 | 2.9 | 3.0 |  |  |
| 2 | 2.6 | 2.7 | 2.5 | 2.8 |  |  |
| 3 | 2.6 | 2.7 | 2.6 | 2.6 |  |  |
| 4 | 2.9 | 2.5 | 2.7 | 2.5 |  |  |
| 5 | 2.9 | 2.6 | 2.5 | 2.6 | 2.7 | 0.818 |

After 32 days, hatchling was converted to fry having average length of 2.7cm. There was no significant difference in final length of hatchling among the ponds (p>0.05).

#### 3.3.3 Weight gain, daily weight gain and specific growth rate of fry

**Table 8.** Initial weight, final weight, weight gain, daily weight gain and SGR of fry during the rearing period of 50 days

| Growth parameters | Pond 1 | Pond 2 | Pond 3 | Pond 4 | Average |
| --- | --- | --- | --- | --- | --- |
| Initial weight(g) | 0.54 | 0.54 | 0.54 | 0.54 | 0.54 |
| Final weight(g) | 21.50 | 21.20 | 20.90 | 20.30 | 20.97 |
| Rearing Period(days) | 50 | 50 | 50 | 50 | 50 |
| Weight gain(g) | 20.96 | 20.66 | 20.36 | 19.76 | 20.43 |
| Daily weight gain(g/fish/day) | 0.42 | 0.41 | 0.41 | 0.40 | 0.41 |
| Specific Growth Rate(%/day) | 7.37 | 7.34 | 7.31 | 7.25 | 7.32 |

The mean weight gain, mean daily weight gain and SGR of fry were found almost similar in all four research pond. Therefore, the mean weight gain, daily weight gain and SGR of fry were 20.43 g, 0.41g/fish/day and 7.32 % per day respectively.

#### 3.3.4 Final length of fry

The final length of fry was measured using scale. After 50 days, fry had attained the size of fingerling. The calculated mean length was tabulated as follow:

**Table 9.**
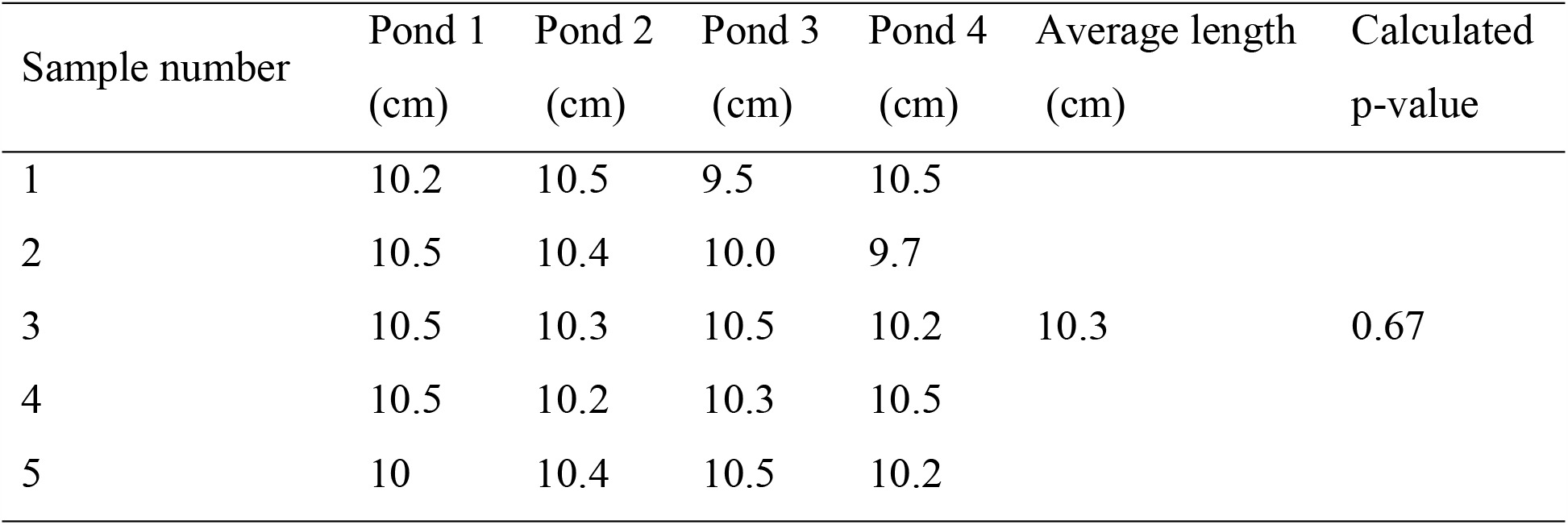
Length of common carp fry in different ponds after rearing for 50 days

After 50 days fry having average length of 2.7cm was converted to fingerling having average length of 10.3cm. There was no significant difference in final length of fry among the ponds (p>0.05).

### 3.4 Survival of fry

Fry mortality was observed from all ponds during the rearing periods. 25,000 fry were stocked in all ponds but on an average 11, 750 fingerling were harvested. Therefore, the survival rate of common carp fry in fish farm of Dhanusha was found to be 47%.

**Table 10.** Number of fry stocked, number of fingerling harvested and survival rate (%) of common carp fry in different ponds

| Pond No. | No. of fry stocked | No. of fingerling harvested | Survival rate (%) |
| --- | --- | --- | --- |
| 1 | 25,000 | 11,500 | 46 |
| 2 | 25,000 | 12,000 | 48 |
| 3 | 25,000 | 11,500 | 46 |
| 4 | 25,000 | 12,000 | 48 |
| Average | 25,000 | 11,750 | 47 |

### 3.5 Water quality

#### 3.5.1 Water quality during breeding and growth of hatchling

The average water temperature in Pond 1,2,3 &4 were 24.8°C, 24.6°C, 25.05°C & 24.75°C respectively during the experimental period of 36 days. The overall water temperature throughout the experiment ranged from 21.8°C-27.6°C. The average water pH in Pond 1, 2, 3 & 4 were 7.9, 7.7, 7.6 & 7.2 respectively. The overall water pH during the experimental period of 36 days ranged from 6.3-8. And the average water DO in Pond 1, 2, 3, & 4 were 5.4 mg/l, 5.3 mg/l, 5.25 mg/l & 5.2 mg/l respectively. The overall water DO during the experimental period ranged from 3.3mg/l −8.9 mg/l.

**Table 11.** Average value and range of water quality parameters during the experimental period of 36 days

| Pond No. | Average value and Range of Water Quality Parameters |  |  |  |  |  |
| --- | --- | --- | --- | --- | --- | --- |
|  | Average temp.<br>(°C) | Range<br>(°C) | Average pH | Range | Average DO<br>(mg/l) | Range<br>(mg/l) |
| 1 | 24.8 | 22.3-27.3 | 7.9 | 7.3-8.4 | 5.4 | 3.7-7.8 |
| 2 | 24.6 | 21.9-27 | 7.7 | 7.3-8.2 | 5.3 | 3.8-7.1 |
| 3 | 25.05 | 22.6-27.6 | 7.6 | 6.8-8.3 | 5.25 | 3.3-8.9 |
| 4 | 24.75 | 21.8-27.6 | 7.2 | 6.3-8 | 5.2 | 4.2-6.6 |

Water temperature, pH & dissolved oxygen in all four ponds were recorded on weekly basis. In each week they were recorded three times a day and their average values were calculated. Calculated average values are tabulated as follow:

**Table 12.**
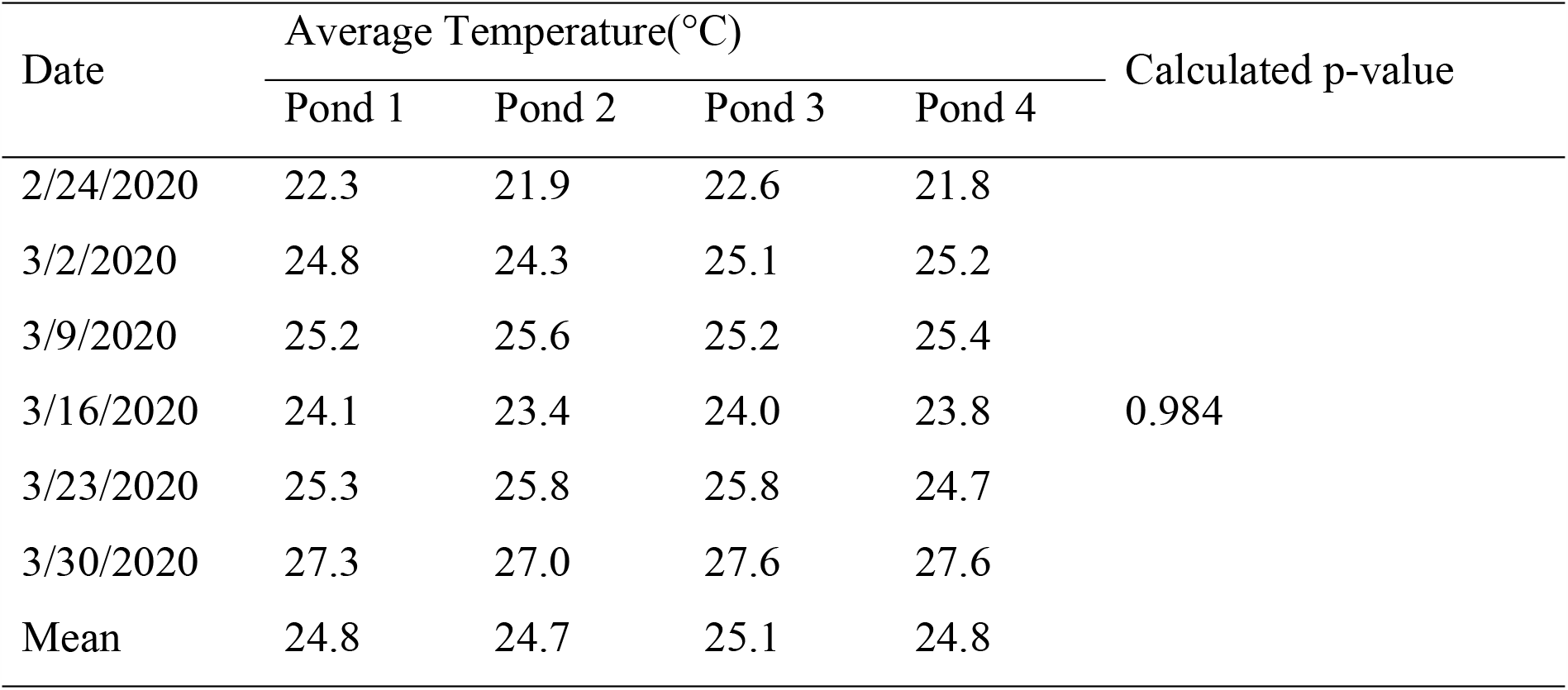
Weekly average water temperature observed in each pond during the experimental period of 36 days

The calculated p value of water temperature is 0.984 which is greater than 0.05. That means there was no significant difference in water temperature among 4 research ponds.

**Table 13.** Weekly average water pH observed in each pond during the experimental period of 36 days

| Date | Average pH |  |  |  | Calculated p-value |
| --- | --- | --- | --- | --- | --- |
|  | Pond 1 | Pond 2 | Pond 3 | Pond 4 |  |
| 2/24/2020 | 7.3 | 7.3 | 6.8 | 6.9 | 0.1 |
| 3/2/2020 | 8.3 | 8.0 | 8.3 | 7.0 |  |
| 3/9/2020 | 8.4 | 8.0 | 8.0 | 8.0 |  |
| 3/16/2020 | 7.7 | 8.2 | 7.7 | 6.3 |  |
| 3/23/2020 | 8.3 | 7.3 | 7.6 | 7.6 |  |
| 3/30/2020 | 7.9 | 7.6 | 7.7 | 7.6 |  |
| Mean | 7.9 | 7.7 | 7.7 | 7.2 |  |

The calculated p value of water pH is 0.10 which is greater than 0.05. That means there was no significant difference in water pH among 4 research ponds.

**Table 14.** Weekly average water DO observed in each pond during the experimental period of 36 days

| Date | Dissolved Oxygen(mg/l) |  |  |  | Calculated p-value |
| --- | --- | --- | --- | --- | --- |
|  | Pond 1 | Pond 2 | Pond 3 | Pond 4 |  |
| 2/24/2020 | 4.4 | 3.8 | 4.5 | 4.2 | 0.984 |
| 3/2/2020 | 4.6 | 4.6 | 5 | 4.9 |  |
| 3/9/2020 | 7.8 | 6.2 | 5.9 | 5.2 |  |
| 3/16/2020 | 3.7 | 4.1 | 3.3 | 4.6 |  |
| 3/23/2020 | 5.8 | 6.1 | 5.9 | 5.7 |  |
| 3/30/2020 | 6.5 | 7.1 | 6.9 | 6.6 |  |
| Mean | 5.47 | 5.32 | 5.25 | 5.20 |  |

The calculated p value of DO is 0.984 which is greater than 0.05. That means there was no significant difference in water DO among four research ponds (p>0.05). and, tabulated values are plotted in graphs as follow:

**Figure 2.**
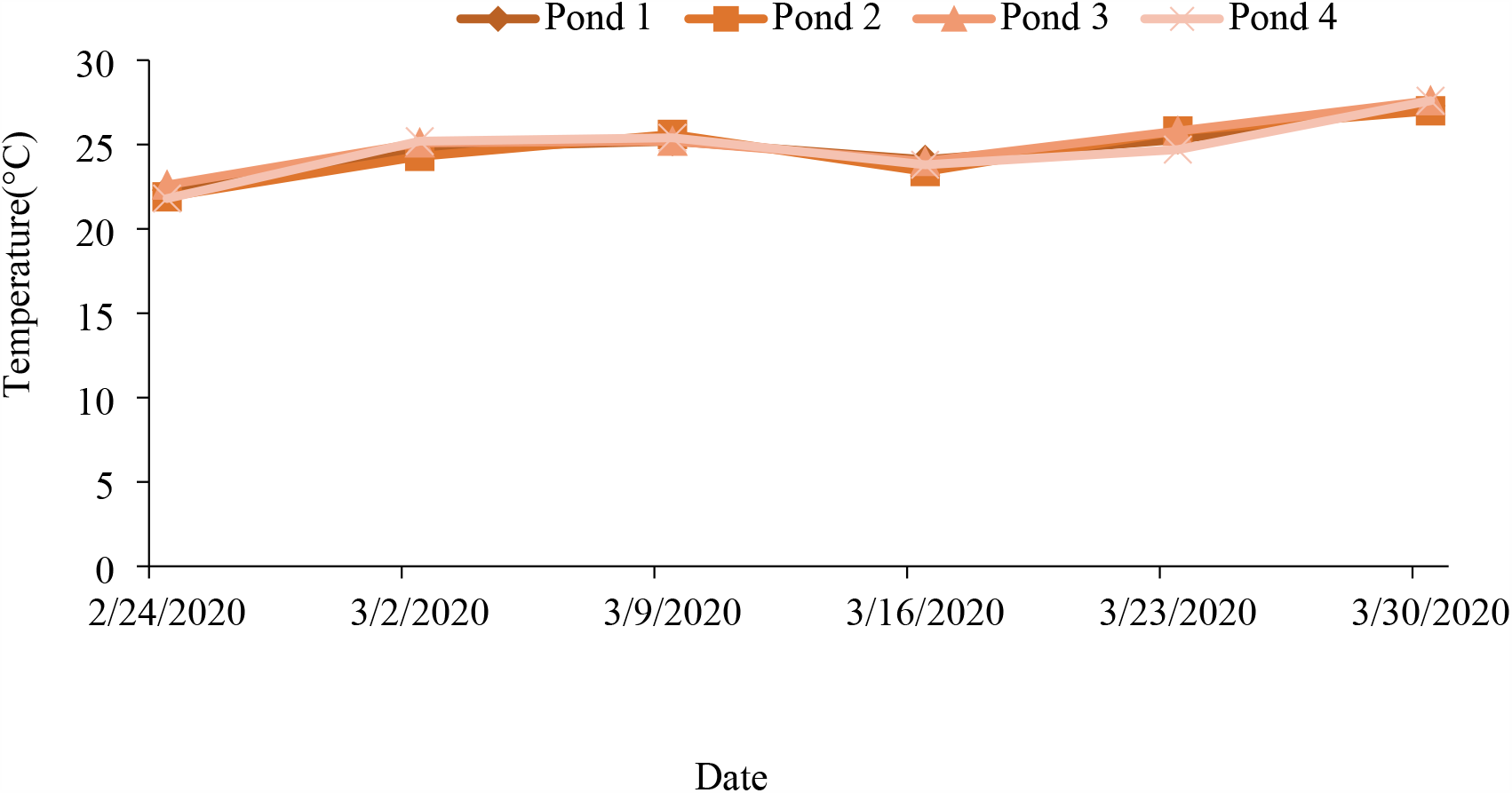
Weekly average water temperature observed in each pond during the experimental.

**Figure 3.**
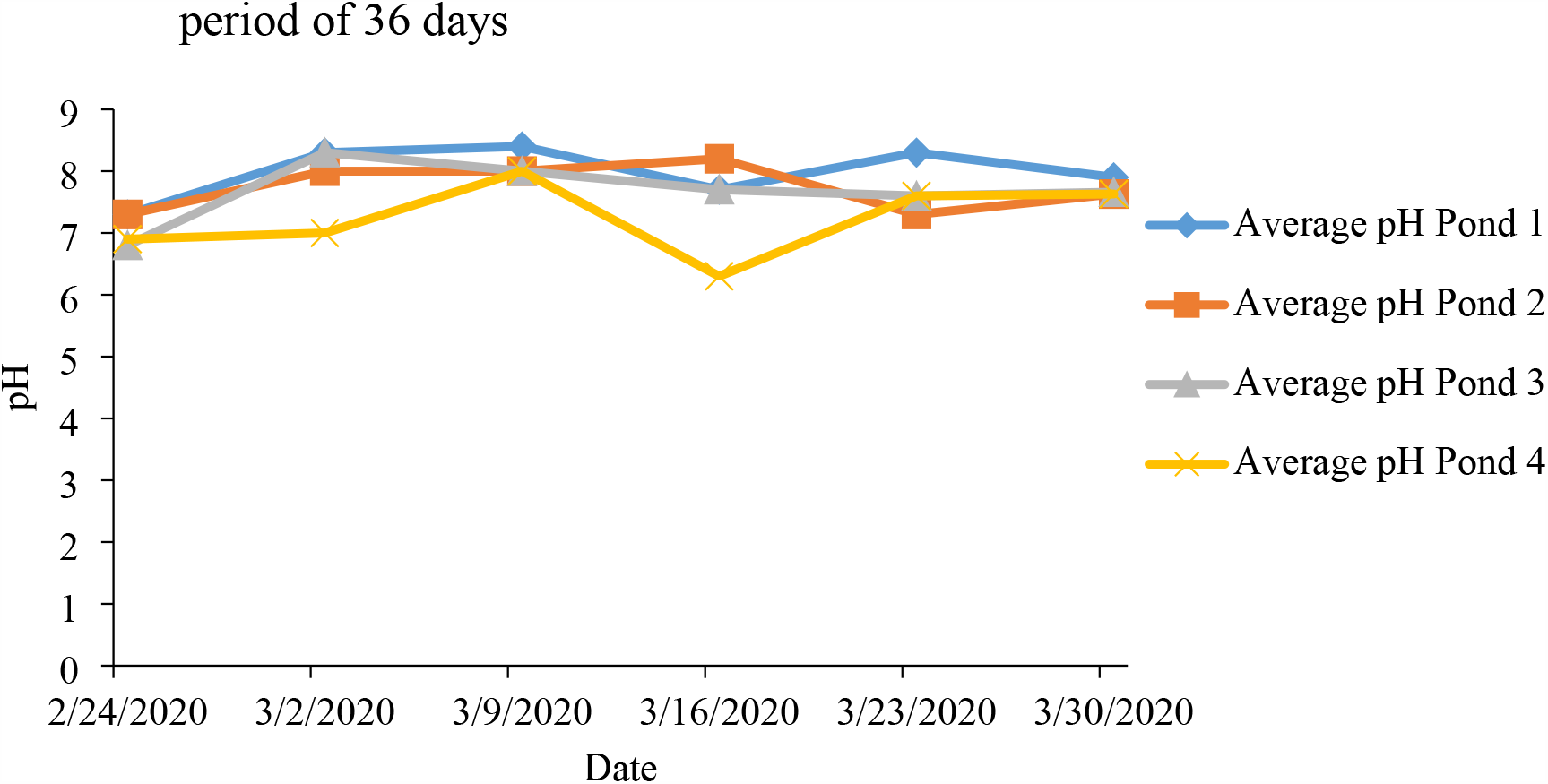
Weekly average water pH observed in each pond during the experimental period of 36 days.

**Figure 4.**
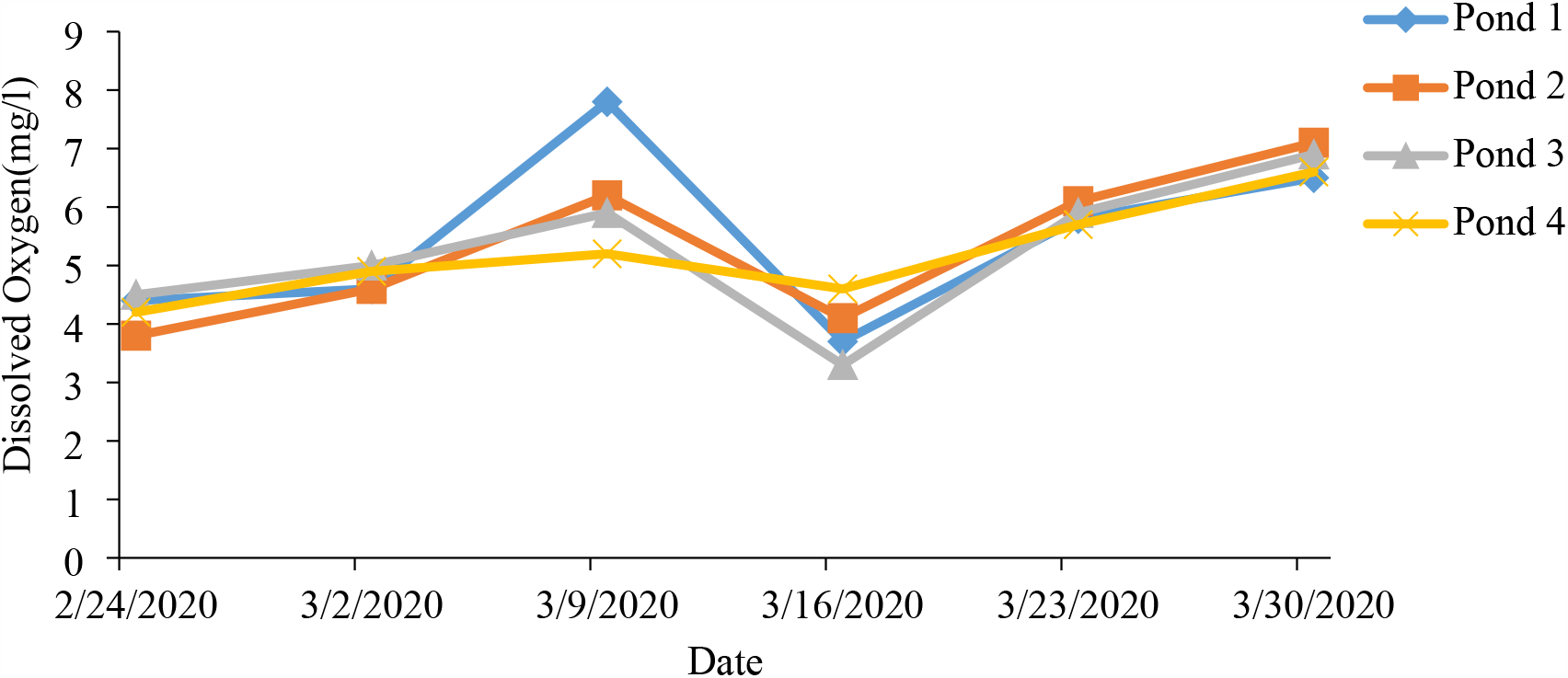
Weekly average water dissolved oxygen observed in each pond during the experimental period of 36 days.

#### 3.5.2 Fluctuation in water quality parameters at different time (morning, day and evening)

Mean temperature of morning, day and evening are tabulated and plotted in graph to observe the fluctuation in water temperature at different time.

**Figure 5.**
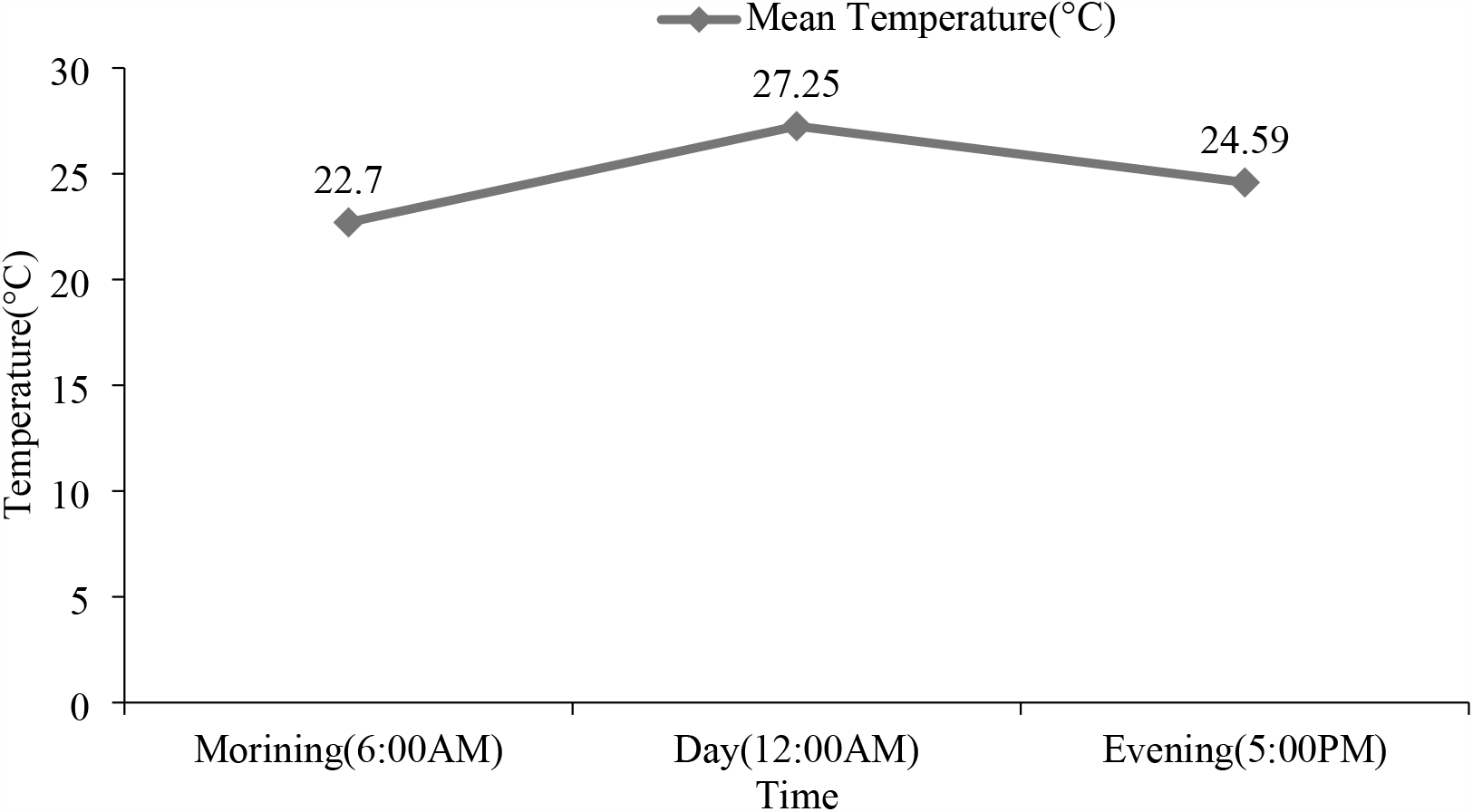
Fluctuation in water temperature observed at different time.

Mean water temperature of morning, day and evening observed during the experimental period of 36 days were 22.7°C, 27.25°C and 24.59°C respectively.

Similarly, mean water pH of morning, day and evening are tabulated and plotted in graph to observe the fluctuation in water pH at different time.

**Figure 6.**
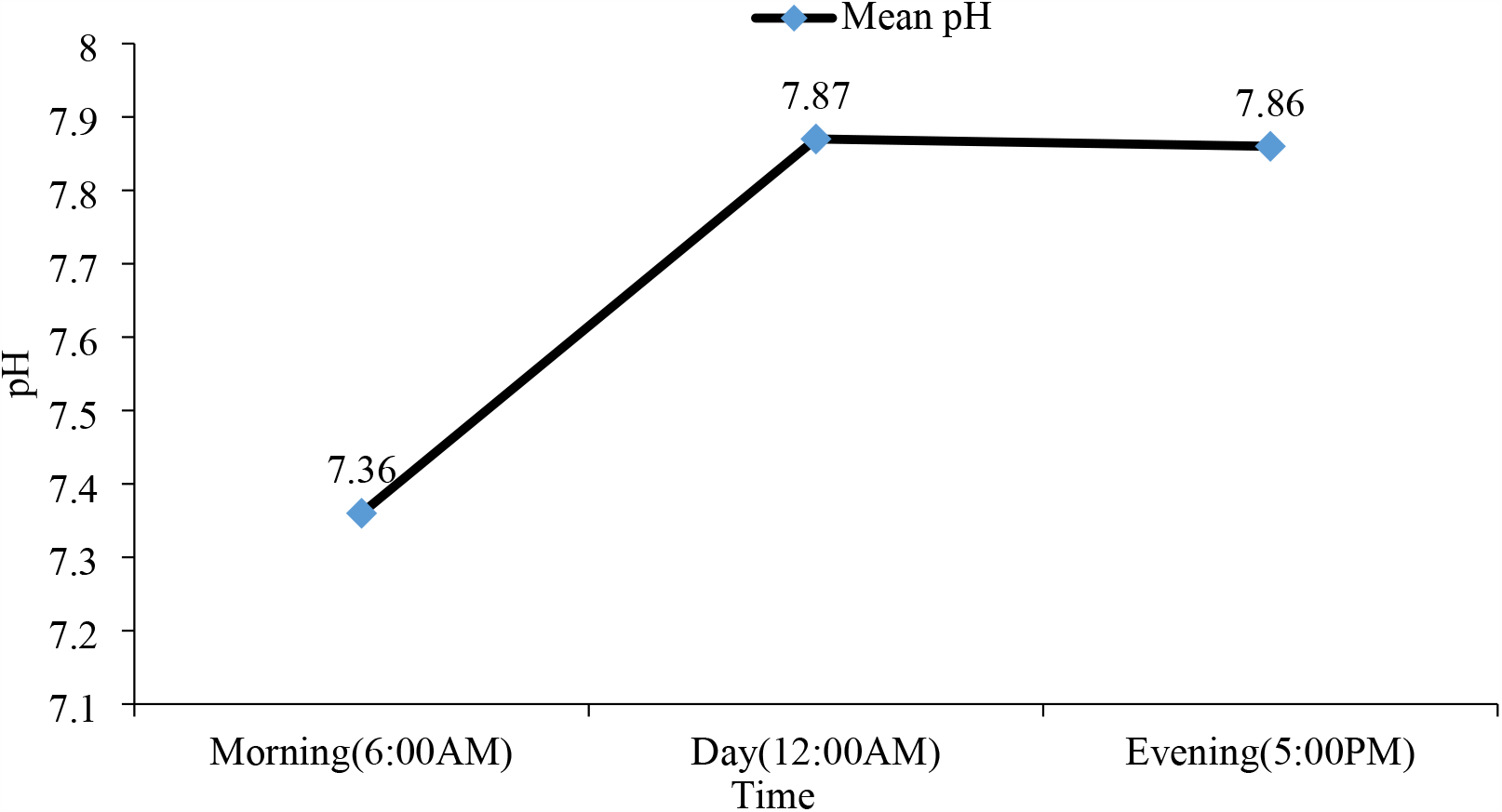
Fluctuation in water pH observed at different time.

Mean water pH of morning, day and evening observed during the experimental period of 36 days were 7.36, 7.87 and 7.86 respectively. No significant fluctuation in water pH was observed at different time.

And, mean water DO of morning, day and evening observed during the experimental period of 36 days are tabulated and plotted in graph to observe the fluctuation in water DO at different time.

**Figure 7.**
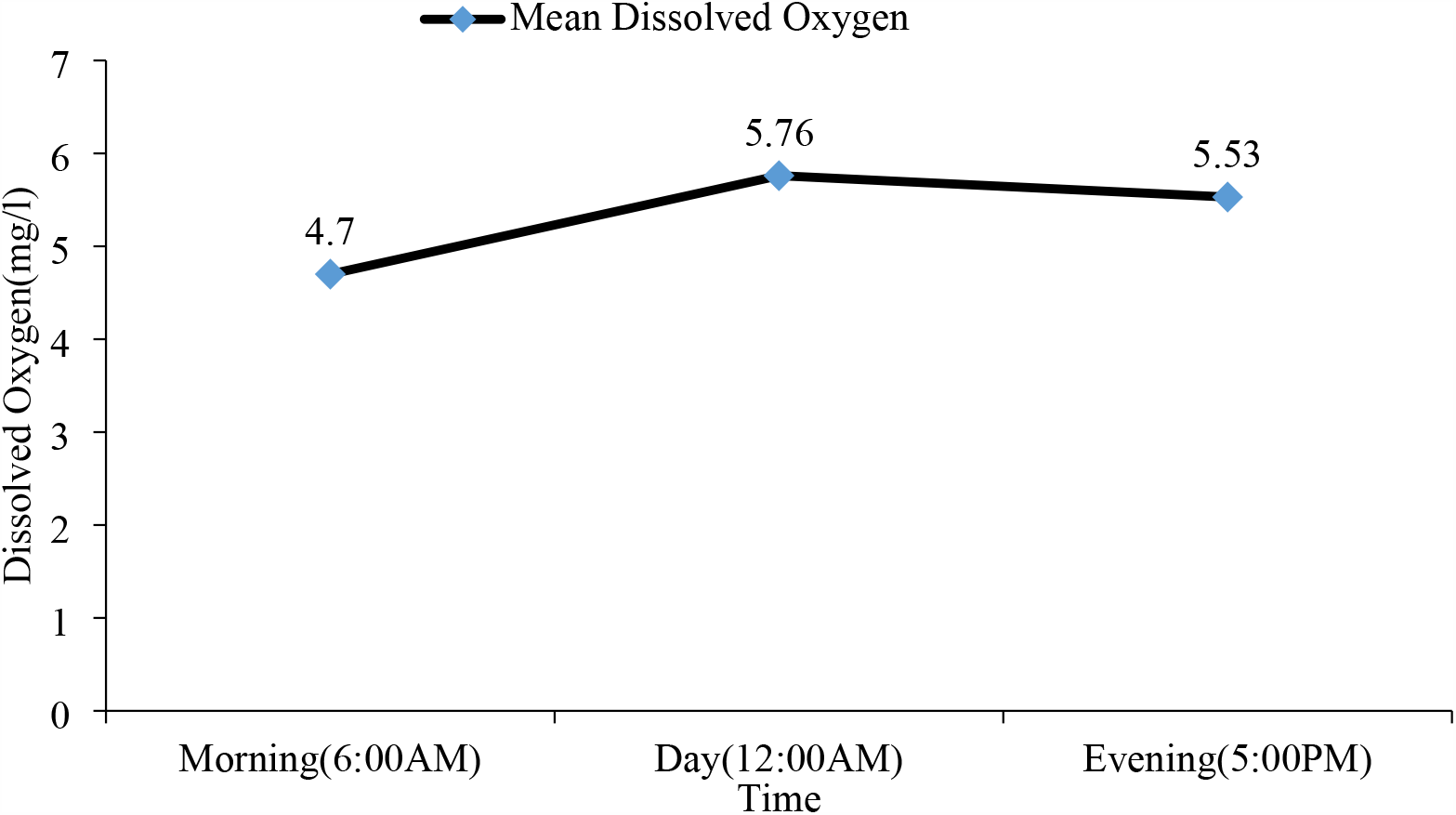
Fluctuation in water DO at different time.

Mean water DO of morning; day and evening observed during the experimental period of 36 days were 4.7mg/l, 5.76mg/l and 5.53 mg/l respectively.

Water temperature, pH and DO were found maximum in day time and were found minimum in morning time.

#### 3.5.3 Water quality during rearing of fry

**Table 15.** Mean value and range of water quality parameters observed during rearing of fry

| Water quality parameters | Range | Average | Calculated p-value |
| --- | --- | --- | --- |
| Temperature (° C) | 27-33 | 30.1 | 0.99 |
| pH | 7-10 | 8.5 | 0.89 |
| Dissolved oxygen (mg/l) | 3-7 | 5.5 | 0.90 |

Calculated p-value for water temperature, pH and dissolved oxygen were greater than 0.05 so there were no significant difference in water temperature, pH and Dissolved oxygen among the ponds during rearing period.

**Figure 8.**
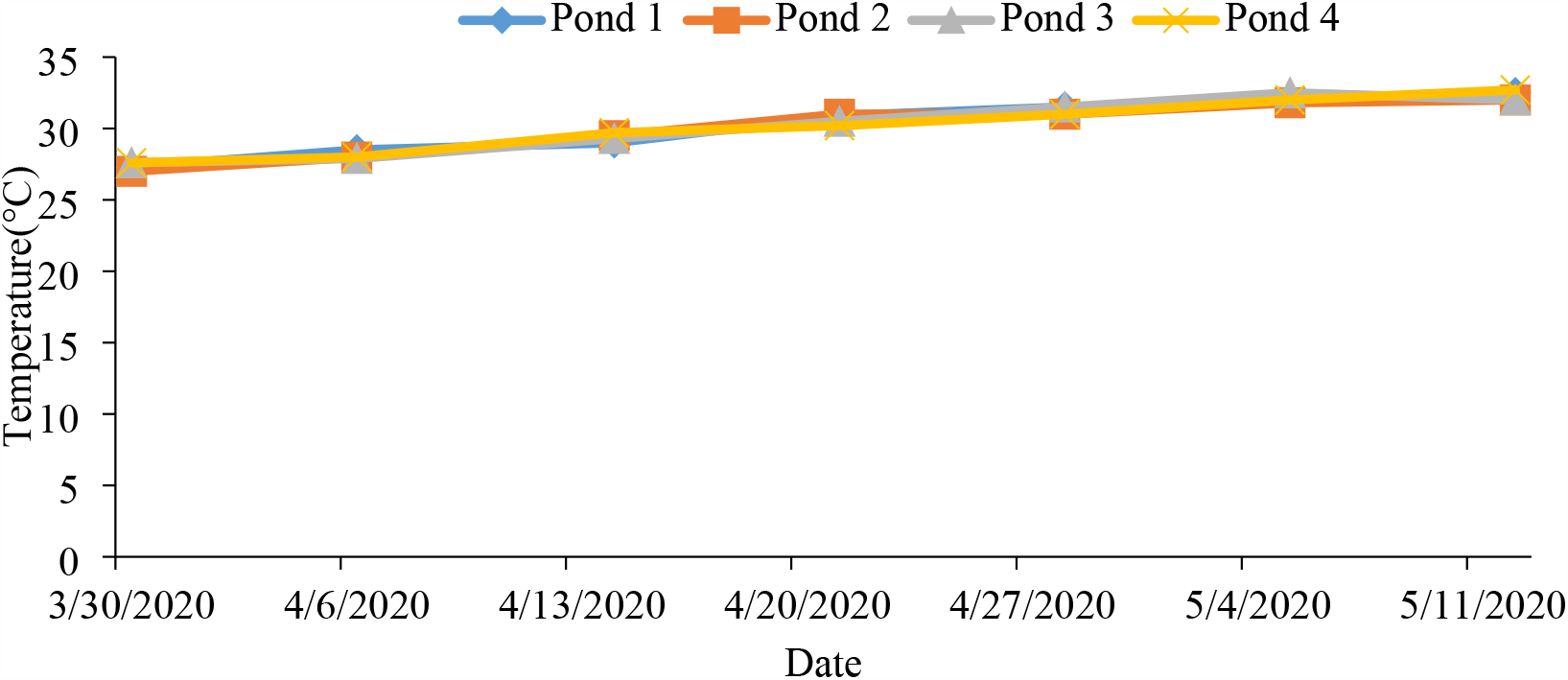
Weekly mean temperature.

**Figure 9.**
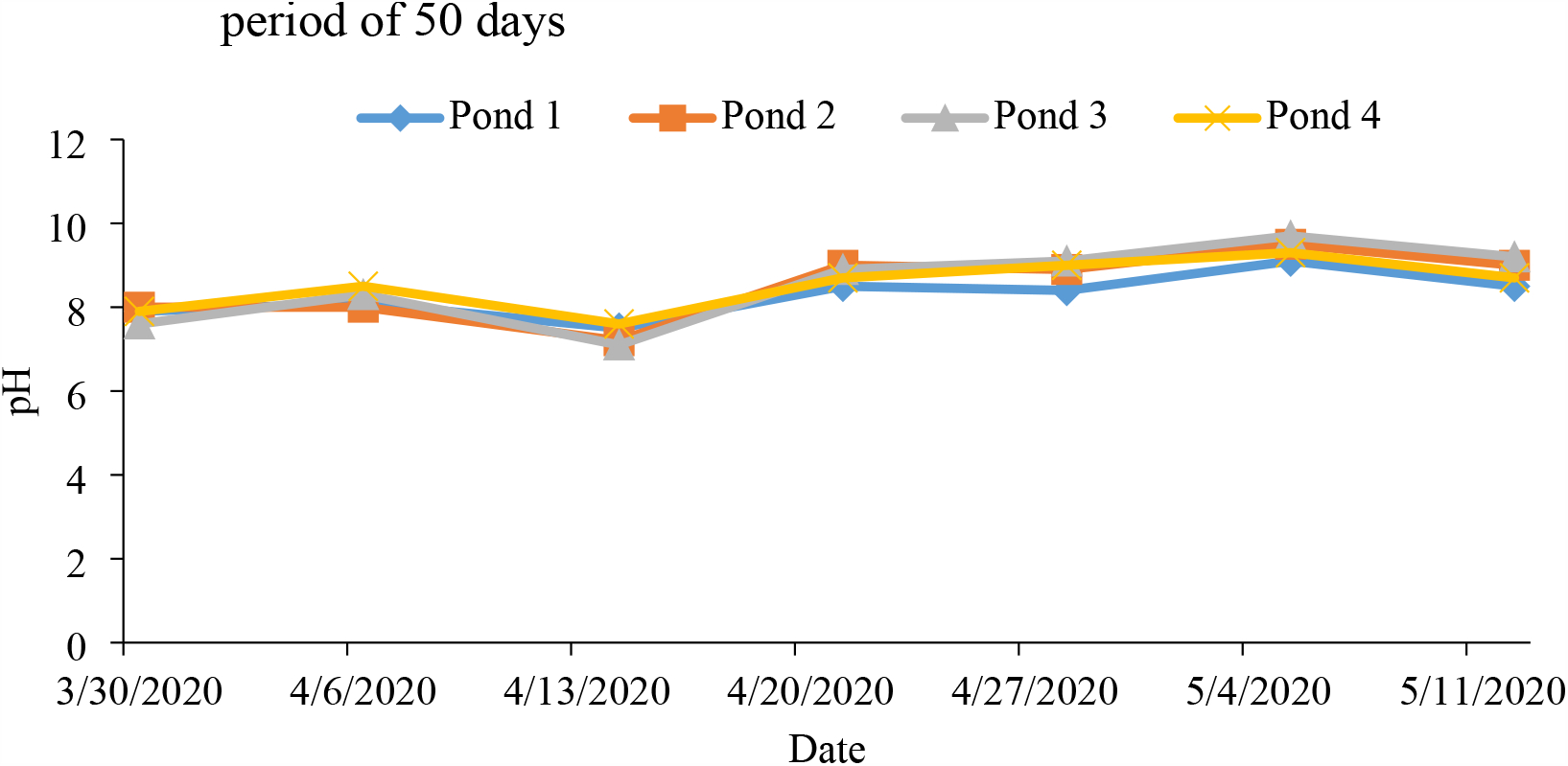
Weekly mean pH of pond water in each pond during the experimental period of 50 days.

**Figure 10.**
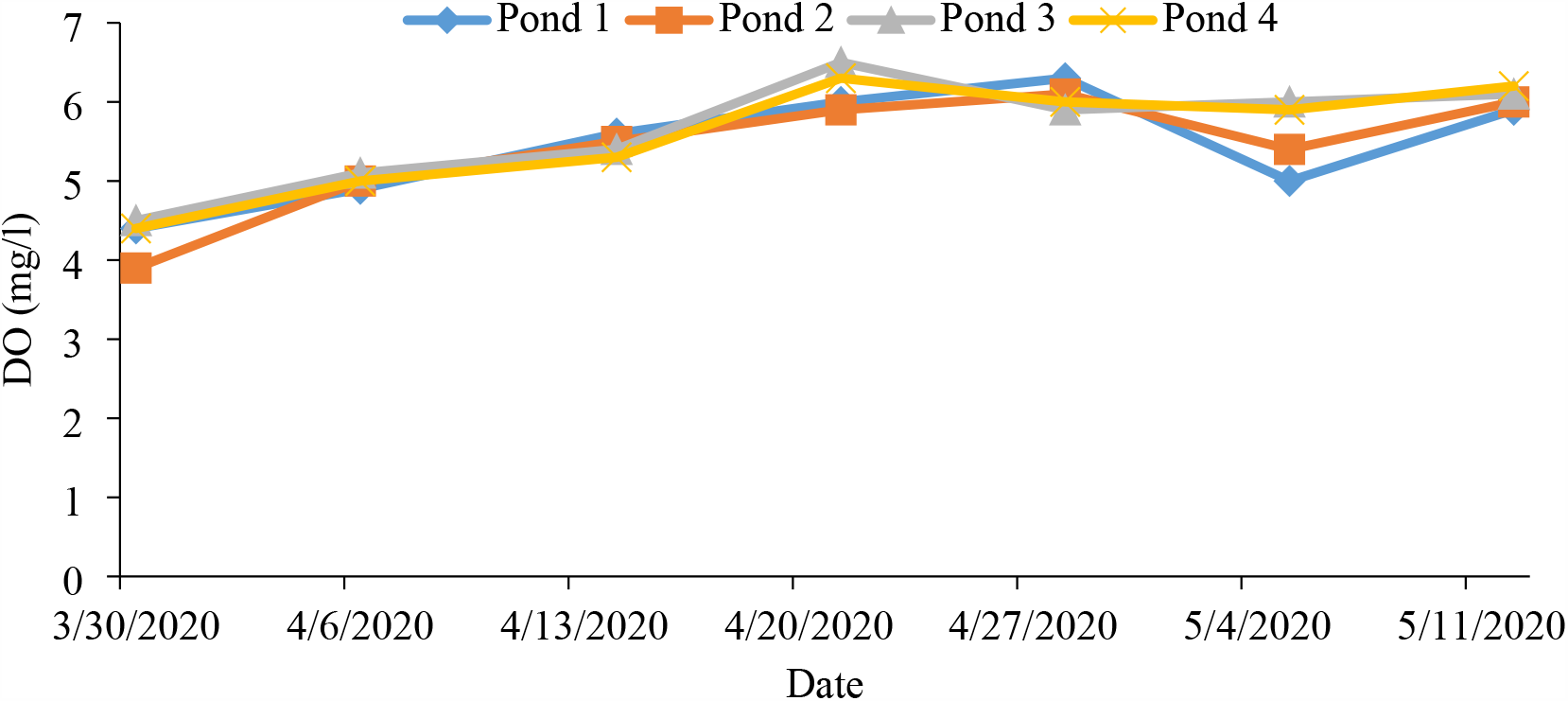
Weekly mean DO of pond water in each pond during the experimental period of 50 days.

## Discussion of fry mortality

Mortality of fry due to predation was high in fish farm of Dhanusha. Predators like king fisher, water snake, white stork, water snake, pond heron etc. were observed. The findings are similar to that showed by Balami and Pokhrel (2020) in their experiment entitled, “Production of common carp and grass carp fingerling in a polyculture system in Chitwan, Nepal”. Predation of fry due to wild fishes like pothiya, tyangra, dhaduwa and gooch was high in fish farm of Dhanusha.

Water temperature and pH recorded during rearing of hatchling and fry were found to be within the suitable ranges as reported by Banjade (2015) where he reported suitable range of water temperature as 20-33°C and suitable range of water pH as 6-9 for better growth of common carp hatchling and fry. Suitable range of water dissolved oxygen for rearing of common carp hatchling and fry as reported by Banjade (2015) is 5.8-9.2 mg/l. But range of water dissolved oxygen was found to be low (i.e. 3-7.8 mg/l) during rearing of hatchling and fry in fish farm of Dhanusha.

Therefore, mortality of common carp hatchling and fry due to predation and low dissolved oxygen was observed in fish farm of Dhanusha.

## 4. SUMMARY, CONCLUSIONS AND SUGGESTIONS

### 4.1 Summary

The study was conducted in Fisheries Human Resource Development and Technology Validation Centre, Janakpurdhaam, Dhanusha. Two consecutive experiments were conducted to assess breeding performance, growth and survival of Common carp. To observe the breeding performance of common carp and growth of its hatchlings, four nursery ponds having same area were selected and research was conducted for 36 days. And, to observe the survival rate of fry, research was conducted in another 4 nursery ponds having same area and research was conducted for 50 days. There were no significant difference in temperature, pH and DO among the ponds (p>0.05) during research period. Feeding schedule and other all conditions were same in all ponds. Hatchlings and fry were fed twice a day. Water quality parameters and weight of hatchling and fry were measured on weekly basis. Since weight of single hatchling is negligible batch weight of hatchling was taken using electronic micro weighing balance.

1,38,750 fry were produced from 7 female having average total weight 19.25kg and 14 male having average total weight 27.77kg through semi-artificial breeding process. The average weight of 4 days old hatchling was 0.02 g. After 29 days the average weight of hatchling was 0.56g. The average weight gain of hatchling was 0.54 g. The daily weight gain and specific growth rate of hatchling were 0.02 g/fish/day and 12.16%/day respectively.Fry were stocked in another 4 ponds at stocking density of 56, 000 fry/ha i.e. 25,000 fry in each pond.

### 4.2 Conclusion

Induced breeding of common carp can be conducted successfully in nursery pond of fish farm, Dhanusha in the month of February when pond water temperature is about 21.8°C. Mortality of fry due to predators and low dissolved oxygen was high in fish farm of Dhanusha. Control of predators, proper water quality management, and quality feed etc. can improve the survival rate of fry in fish farm of Dhanusha. The stocking weight of fry was 0.54g. After 50 days the average final weight of fry was 20.97g. The average weight gain of fry was 20.43g. The daily weight gain and specific growth rate of fry were 0.41 g/fish/day and 7.32%/day respectively. The average survival rate of fry at stocking density of 56,000/ha was 47%. Mortality of fry due to predators like king fisher (*Alcedo atthis*), water snake (*Neordia sipedon*), white stork (*Ciconia ciconia*), pond heron (*Ardeola grayii*) was high in fish farm of Dhanusha. Mortality of fry due to wild fises like pothiya, tyangra, dhaduwa was also high. Mortality of fry during rearing period might be due to asphyxiation because water Dissolved Oxygen was low during.

